# Talk2Biomodels: AI agent-based open-source LLM initiative for kinetic biological models

**DOI:** 10.1101/2025.03.11.642548

**Authors:** Lilija Wehling, Gurdeep Singh, Ahmad Wisnu Mulyadi, Rakesh Hadne Sreenath, Henning Hermjakob, Tung Nguyen, Thomas Rückle, Mohammed H. Mosa, Henrik Cordes, Tommaso Andreani, Thomas Klabunde, Rahuman S. Malik-Sheriff, Douglas McCloskey

## Abstract

In this study, we present Talk2Biomodels (T2B), an open-source^1^, user-friendly, large language model-based agentic AI platform designed to democratize access to computational models of biological interactions and promote the FAIRification (Findability, Accessibility, Interoperability, and Reusability) of these models. T2B enables users to explore and analyse mathematical models of biological systems using natural language. It eschews the traditional graphical user interface (GUI) and minimally adaptable workflow in favour of a modern agentic framework to provide a dynamic and immersive experience to explore mathematical models of biological interactions through conversations in natural language. T2B supports models encoded in the open-source community format Systems Biology Markup Language (SBML) and is integrated with the BioModels database (https://www.ebi.ac.uk/biomodels/), enabling seamless exploration, simulation, and analysis of curated systems biology models. Use cases in precision medicine, epidemiology, and emergent systems properties of biological networks are presented to demonstrate how experts and non-experts in computational biology can benefit from T2B.

## Introduction

Recent developments in generative artificial intelligence (AI), particularly in large language models (LLMs), have demonstrated substantial progress. Noteworthy advances in both algorithmic innovation and data-driven biomedical discovery are showcased by the advent of foundational learning models for protein structure, single cell omics and genomic data across species, such as, AlphaFold, scGPT, and Evo 2, to name a few (Brixi et al., 2025, p. 2; Cui et al., 2024; Jumper et al., 2021). While LLMs’ role is currently confined to narrow, task-specific applications, efforts are underway to enable open-source LLM-driven agentic workflows in biomedical discovery (Lobentanzer et al., 2025). Collaborative AI agents, with their task orchestration, reflective learning and sophisticated reasoning capabilities, have the potential to revolutionize biomedical research and patient treatment (Gao et al., 2024; Zou & Topol, 2025).

Quantitative computational models have become an essential component in enhancing our understanding of biological systems. Besides the qualitative knowledge of biological interactions, they promote an engineering perspective by enforcing quantification of systems properties and their interactions. Thus they enable scientists to predict the outcomes of perturbations in complex biological networks and serve as analytical tools to test the theoretical understanding of the dynamics within biological systems (Saez-Rodriguez & Blüthgen, 2020).

The most common frameworks in systems biology modeling include kinetic, constraint-based, logic, and agent-based modeling. The present study specifically focuses on ODE models, a subset of kinetic models, due to their widespread application and significance in the field. ODE models represent the most popular modeling approach, with approximately 1662 models^2^ currently deposited in the BioModels database (Malik-Sheriff et al., 2020) and a multitude not systematically stored models in scientific literature. These models cover a wide range of domains, as demonstrated by their use in precision medicine (e.g., inflammatory bowel disease treatment) (Dwivedi et al., 2014), COVID-19 pandemic studies (Tang et al., 2020), and theoretical analysis of dynamic systems (Markevich et al., 2004). Furthermore, ODEs are a standard tool in pharmaceutical research, particularly in the development of physiologically based pharmacokinetic/pharmacodynamic (PBPK/PD) and quantitative systems pharmacology and toxicology (QSP/QST) models (Chan et al., 2024; Kuepfer et al., 2016). In addition, the regulatory authorities, such as the U.S. Food and Drug Administration (FDA), recognize these computational models as a well-established and increasingly significant body of evidence in drug approval decision-making processes (Morrison et al., 2018).

To interact with mathematical models of biological systems, several popular graphical user interface (GUI)-based ordinary differential equations (ODE) modelling frameworks exist for biological and translational applications. These include proprietary options such as the MATLAB toolbox SimBiology (The MathWorks Inc., 2024) and Berkley Madonna (Marcoline et al., 2022), as well as open-source alternatives like COPASI (Hoops et al., 2006), the Open Systems Pharmacology Suite (Kuepfer et al., 2016) and RunBioSimulations (Shaikh et al., 2021). However, both proprietary and open-source solutions typically require software installation, specialized training, and domain knowledge for effective simulation and analysis of models.

To overcome the limitations of existing modelling frameworks, we propose Talk2BioModels (T2B), an easily accessible chatbot-based ODE modelling platform built using agentic AI. T2B users are not required to install software or scripting frameworks, nor do they need to master programming languages. Additionally, T2B users do not need to possess expert knowledge in systems biology to simulate and perform basic analyses of ODE models. The platform supports the import and export of models encoded in SBML (Keating et al., 2020) and ensures interoperability, independent of the operating system or programming framework. T2B also provides access to the BioModels database, which hosts the largest collection of curated quantitative mathematical models (Malik-Sheriff et al., 2020).

By facilitating the sharing and simulation of mathematical models in SBML format, T2B improves communication between computational scientists and experimental biologists, enhances reproducibility of experimental findings, and enables efficient searching and reuse of models, thus contributing to the FAIR principles (Findable, Accessible, Interoperable, and Reusable) of data exchange (Wilkinson et al., 2016). It extends the application of mathematical models to other research fields, such as medicine, and offers an engaging environment for teaching and exploring a variety of models. Moreover, T2B empowers non-experts to interact with models and pose simulation-related questions directly in natural language, aided by LLMs grounded on model simulations for interpreting the data without hallucination.

## Materials and Methods

We developed the T2B application using the LangGraph library (v0.3.2)^3^, which provides a framework for constructing stateful computational graphs. In this architecture, each node represents a discrete computational step, while the edges define the relationships between these steps. The application maintains a global state - a snapshot of the application - which is propagated and updated throughout the execution process (Fig 1).

**Figure 1:**
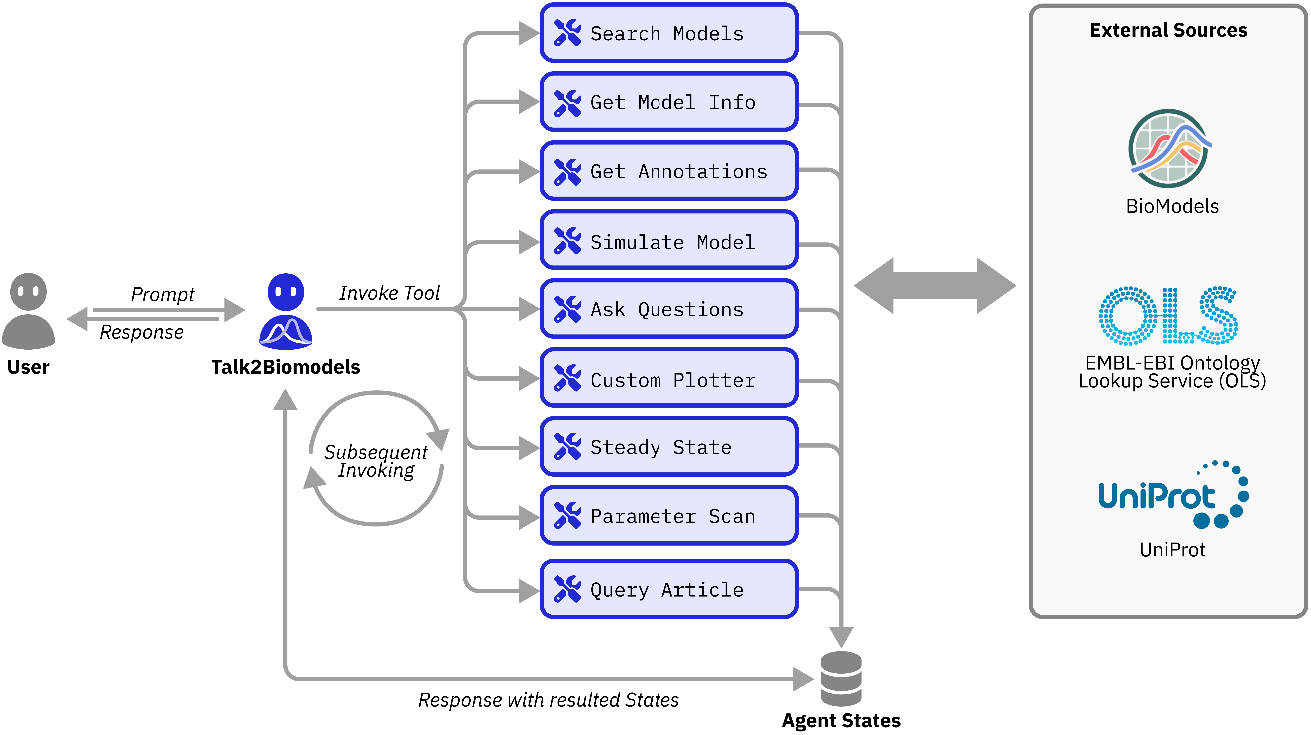
Architecture diagram of Talk2Biomodels. Users can prompt the T2B agent to invoke various tools for model search, simulation and plotting, stability analysis, annotation extraction and retrieval-augmented generation (RAG) on uploaded scientific articles. These agentic AI tools interactively engage with external resources, including BioModels, OLS, and UniProt databases. T2B maintains the global state of the agent throughout the chat history. BioModel search, simulations, annotation extraction and stability analysis are carried out using BASICO python library.

In the case of T2B, the computational nodes consist of a prebuilt reasoning and action (ReAct)- based agent, implemented following the ReAct paradigm (Yao et al., 2023), and a suite of tools available to the agent (Fig 1). Specialized operation on the models are carried out using the BASICO library (v0.78) (Bergmann, 2023). Directed edges in the graph structure facilitate interactions between the agent and these tools. The agent autonomously selects and executes tool invocations based on user input, dynamically updating the global state during execution. The state management mechanism tracks key-value pairs storing messages, analysis outputs, and intermediate results, ensuring contextual coherence throughout the workflow. Upon computation completion, the last updated state is parsed to extract relevant information and render it for the user.

For configuring agent-based text generation tasks, users can select between two LLMs: OpenAI’s GPT-4o-mini (OpenAI et al., 2024) and NVIDIA NIM-optimized Meta/Llama-3.3-70B- Instruct^4^. For text-to-embedding transformations, the application supports two embedding models: OpenAI’s text-embedding-ada-002 and NVIDIA NIM-optimized Llama-3.2-NV- EmbedQA-1B-V2^5^. The backend of T2B is implemented in Python 3.12, while the front end was developed using Streamlit^6^ (v1.39.0) and Plotly^7^ (v5.24.1) for an interactive and user-friendly interface.

The T2B application is hosted on GitHub (https://github.com/VirtualPatientEngine/AIAgents4Pharma), facilitating open-source contributions and discussions, and is linked to individual models at BioModels repository.

## Results and Discussion

T2B offers a comprehensive suite of functions for biological and translational ODE models. Users can upload ODE models in SBML format or retrieve curated models from the BioModels database using the *search models* tool (Fig 2). The *get model information* tool extracts model details, including species, parameters, and model abstract. Concurrently, the *get annotations* tool retrieves and displays model species’ identifiers by dynamically accessing Ontology Lookup Service (OLS) and UniProt data bases (Jupp et al., 2015; The UniProt Consortium, 2025) over APIs. This capability enables researchers to thoroughly explore and interpret the specific nomenclature of the model.

**Figure 2:**
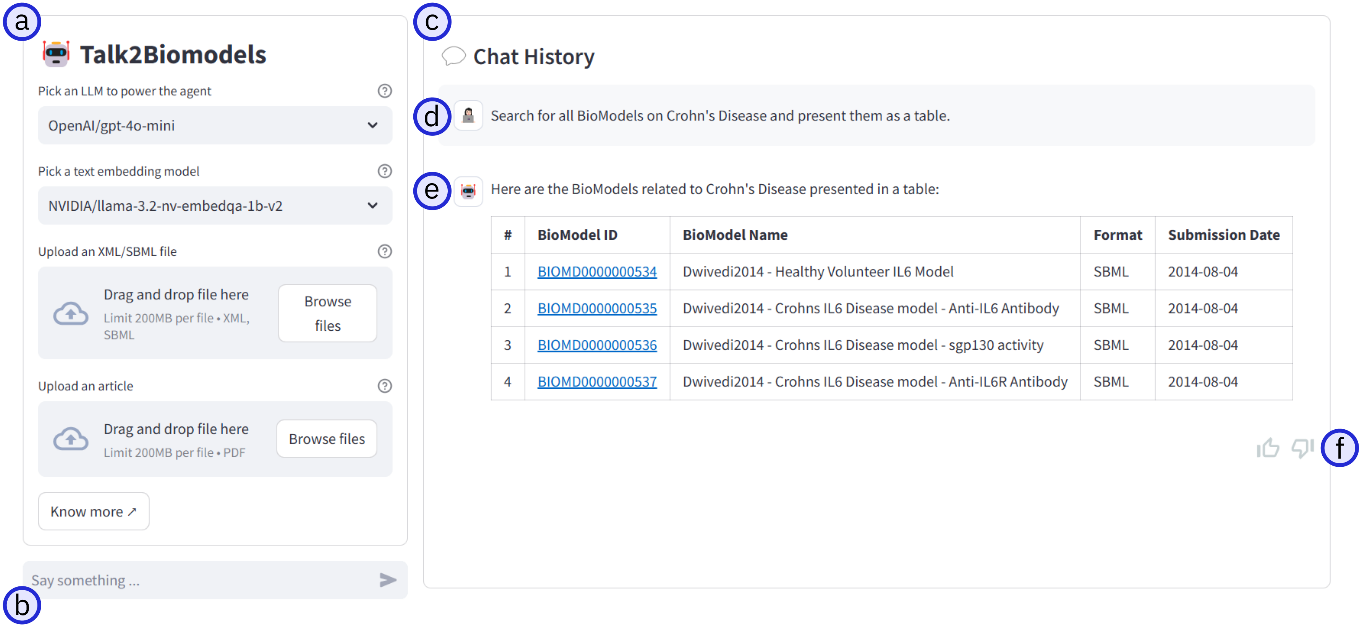
An example of the search models tool in the Talk2BioModels GUI. This GUI supports both open and closed-source LLMs, enabling users to upload SBML-formatmathematical models and PDF-format scientific articles (a). T2B allows natural language prompts via the chat box interface (b). The GUI features a chat history window (c) displaying question-and-answer conversations (d, e). Additionally, T2B’s GUI allows dynamic user feedback through upvote and downvote buttons. (f). Here, we exemplify *search models* tool for keyword-based searches in the BioModels database (e).

In T2B, SBML models can be directly accessed, simulated, and analyzed. The platform supports time-course simulations with adjustable time steps, initial concentrations and durations (*simulate model* tool), presenting results in dynamic tables and graphs that can be interactively modified to suit the users’ needs (*custom plotter* tool). Additionally, users can interrogate model description and simulation results (*ask questions* tool) and alter model parameters using natural language.

Moreover, T2B offers several tools for model analysis: the *steady state* tool determines whether a model reaches equilibrium, while the *parameter scan* tool performs multiple simulations by systematically varying model initial conditions or parameter values within user-specified range and intervals (Fig 3). Finally, the *query article* tool leverages retrieval-augmented generation (RAG) framework (Lewis et al., 2020) to extract supplementary information from user-uploaded research articles, enhancing model understanding.

**Figure 3:**
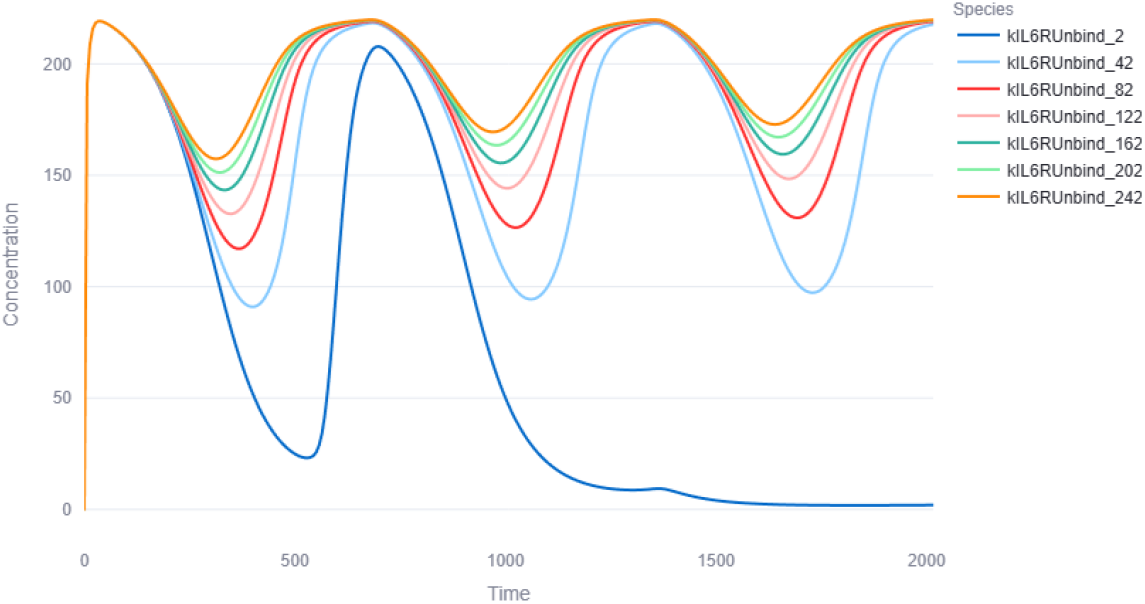
Parameter scan of a precision medicine model (Dwivedi et al., 2014) within T2B GUI. A parameter ‘kIL6Unbind’ is varied in steps of 40 units from 2 to 242 to simulate an unbinding event of an antibody from its target interleukin 6 receptor (IL6R). Each curve represents a separate simulation of this model.

Overall, this suite of agentic AI-driven tools offers a unique platform for working with ODE models, combining user-friendliness and interactive features.

### Use-cases

Use cases in precision medicine^8^, epidemiology^9^, and emergent systems properties^10^ of biological networks are presented below and made available on GitHub to demonstrate how experts and non-experts in computational biology can benefit from T2B. All three examples illustrate the processes of loading, simulating, interactively plotting, and editing models, as well as how LLMs assist in drawing conclusions from the model simulations.

#### Precision Medicine

This example utilizes a multiscale model of interleukin 6 (IL-6) mediated immune regulation in Crohn’s disease to examine drug-disease interaction mechanisms and drug treatment prediction (Dwivedi et al., 2014). The model was validated using clinical trial biomarker data from patients treated with tocilizumab, an anti IL6 receptor monoclonal antibody drug.

Users can interrogate model annotations, adjust the dosage of a drug, simulate drug administration at intervals of either 2 or 4 weeks, and view the value of the clinical readout parameter (CRP in serum) at any given time point. Additionally, they can determine the time required for the depletion of serum anti-IL6Ralpha concentration by replicating the published figure (Dwivedi et al., 2014). The effect on CRP suppression can also be compared by simulating different antibodies with varying dissociation affinities, thereby performing *in silico* tests for potential antibodies.

#### Epidemiology

This example utilizes an epidemiological model to predict the spread of COVID- 19 (Tang et al., 2020). T2B users can predict whether quarantine as a COVID-19 safety measures will help to contain pandemics and predict the impact of the quarantine rate on the number of infected individuals over time, thereby assessing the effectiveness of quarantine measures.

#### Emergent properties of biological networks

This example explores intracellular signaling of the MAP kinase pathway (Markevich et al., 2004) to illustrate emergent behaviors such as hysteresis and bistability induced by covalent modifications of proteins, which serve as widespread regulatory mechanisms in living systems. T2B users can better understand the description of model components and demonstrate hysteresis by performing steady-state analysis for two different stable states.

## Conclusions

T2B is an open-source, user-friendly agentic AI platform designed to democratize access to computational models of biological interactions. By providing a natural language interface, supporting open-source and community standards, and integrating with BioModels data base, T2B reduces the barrier for non-experts to benefit from computational biology models, facilitates experts to disseminate systems biology, and overall significantly contributes to the FAIRification (by improving Findability, Accessibility, Interoperability, and Reusability) of models (Wilkinson et al., 2016). User cases in precision medicine, epidemiology, and the exploration of emergent system properties were presented to demonstrate the ease at which T2B users can reproduce and explore mathematical modelling results.

With an expanding user base of T2B and the increasing complexity of agentic tools and workflows, as well as real-life decisions based on model outcomes, it is crucial to ensure transparency and validation of results. Minimizing hallucinations through model simulation-specific tool calling is essential and must be rigorously tested using comprehensive testing suites. Additionally, as new LLMs are developed and incorporated into the T2B framework, it is important to continuously assess and validate their reasoning capabilities.

While T2B is a significant step in democratizing access to systems biology models, there are several limitations that future work will need to address. First, most mathematical models used in industry are not built using open-source frameworks. We aim to introduce features that will support the translation and reannotation of mathematical models according to open-source and community standards. Second, better model reusability beyond replication of existing results requires refitting of model parameters to new datasets. We aim to add features that will facilitate the mapping of data onto mathematical models and the refitting of model parameters to new datasets. Third, it is essential to verify that user input values for initial concentrations or parameter values are within a reasonable range suitable for the biological system being modeled. LLMs can assist by suggesting appropriate biological ranges and substantiating these suggestions with references from scientific publications. Fourth, expanding the modeling frameworks beyond ODEs is necessary. We would greatly appreciate contributions to our open-source project from the community.

## Acknowledgements

We acknowledge the contributions of Anil Kumar Kanasani and Maryam Najafian during the “AI Agents for Life Sciences” hackathon on October 14-15 2024. We acknowledge the technical support from Dror Hilman, Jake Valsamis, Conor Mohan, and Ben Evan Tsur, gracious donations of computing resources during our hackathon from Shahar Frunkin, and Daniel Koster, and an going support and facilitation from Simon Adar, all from CodeOcean https://codeocean.com/. We acknowledge funding support for our Hackathons from the AI Health Innovation Cluster https://www.aih-cluster.ai/ and particular the help received from Daniela Beyer and the rest of the AIH Coordination Team. We thank Ornella Kossi, Ann-Kristin Mueller, and Stefanie Schimmel from BioLabs Heidelberg, https://www.biolabs.io/heidelberg, for hosting our hackathons. We thank Isabel Wilkinson, Uwe Samer, Nick Venanzi, and David Ruau from NVIDIA for assisting with our integration of NVIDIA NIMs for chat and embedding inference and graciously providing free credits for use with T2B. We thank Vultr https://www.vultr.com/ for graciously providing free credits for deploying T2B.

## Author contributions

LW, GS and DM conceptualized the study. GS developed the agentic AI framework and the agentic AI tools for T2B. AWM contributed to the development of the agentic AI framework and created the figures and icons. RHS implemented *get annotations* tool. GS and AWM developed T2B’s user interface. HH and TN crosslinked T2B to individual models in BioModels repository. DM, RSMS, and LW conceptualized use cases. LW wrote the manuscript. DM supervised the writing of the manuscript and the development of the agentic AI framework. All authors contributed to testing T2B and reviewing the manuscript.

## Footnotes

1 Source code available at: https://github.com/VirtualPatientEngine/AIAgents4Pharma

2 Viewed on 24.02.2025

3 (https://github.com/langchain-ai/langgraph

4 https://build.nvidia.com/meta/llama-3_3-70b-instruct/modelcard

5 https://build.nvidia.com/nvidia/llama-3_2-nv-embedqa-1b-v2

6 https://streamlit.io/

7 https://plotly.com/chart-studio-help/citations/

8 https://virtualpatientengine.github.io/AIAgents4Pharma/talk2biomodels/cases/Case_1/

9 https://virtualpatientengine.github.io/AIAgents4Pharma/talk2biomodels/cases/Case_2/

10 https://virtualpatientengine.github.io/AIAgents4Pharma/talk2biomodels/cases/Case_3/

